# Exploring Genetic Variation that Influences Brain Methylation in Attention-Deficit/Hyperactivity Disorder

**DOI:** 10.1101/413005

**Authors:** Laura Pineda-Cirera, Anu Shivalikanjli, Judit Cabana-Domínguez, Ditte Demontis, Veera M. Rajagopal, Anders D Børglum, Stephen V. Faraone, Bru Cormand, Noèlia Fernàndez-Castillo

## Abstract

Attention-deficit/hyperactivity disorder (ADHD) is a neurodevelopmental disorder caused by an interplay of genetic and environmental factors. Epigenetics is crucial to lasting changes in gene expression in the brain. Recent studies suggest a role for DNA methylation in ADHD. We explored the contribution to ADHD of allele-specific methylation (ASM), an epigenetic mechanism that involves SNPs correlating with differential levels of DNA methylation at CpG sites. We selected 3,896 tagSNPs reported to influence methylation in human brain regions and performed a case-control association study using the summary statistics from the largest GWAS meta-analysis of ADHD, comprising 20,183 cases and 35,191 controls. We identified associations with eight tagSNPs that were significant at a 5% False Discovery Rate (FDR). These SNPs correlated with methylation of CpG sites lying in the promoter regions of six genes. Since methylation may affect gene expression, we inspected these ASM SNPs together with 52 ASM SNPs in high LD with them for eQTLs in brain tissues and observed that the expression of three of those genes was affected by them. ADHD risk alleles correlated with increased expression (and decreased methylation) of *ARTN* and *PIDD1* and with a decreased expression (and increased methylation) of *C2orf82*. Furthermore, these three genes were predicted to have altered expression in ADHD, and genetic variants in *C2orf82* correlated with brain volumes. In summary, we followed a systematic approach to identify risk variants for ADHD that correlated with differential cis-methylation, identifying three novel genes contributing to the disorder.

## INTRODUCTION

Attention-deficit/hyperactivity disorder (ADHD) is a common neurodevelopmental disorder with a worldwide prevalence of around 5% ^1^. Its main symptoms include inattention and/or hyperactivity-impulsivity (DSM-V) ^2^. ADHD is among the most heritable psychiatric disorders, with about 76% of its etiology accounted for by genetic risk factors ^3^ and with single-nucleotide polymorphisms (SNPs) explaining around 22% of the phenotypic variance ^4^. Furthermore, there is molecular evidence of shared genetic risk factors across the main five psychiatric disorders (ADHD, autism spectrum disorders (ASD), schizophrenia, bipolar disorder (BD) and major depression disorder (MDD)) ^5^. In ADHD, a recent genome-wide association study meta-analysis of 12 sample groups unraveled some of the specific genetic underpinnings of this polygenic disorder for the first time ^4^. One of the challenges of genome-wide association studies (GWAS) is to establish the causal relationship between the associated genetic variants, especially those located outside genes, and the disorder. In this regard, the use of epigenetic information can improve the interpretation of functionality of non-coding genetic variation ^5,6^. In addition, some studies have hypothesized the importance of sub-threshold variants derived from GWAS ^7^, particularly those located in enhancer regions, with a potential impact on gene regulation ^8,9^.

DNA methylation is one of the most stable epigenetic mechanisms, involving methylation mainly at cytosines of CpG dinucleotides. This mechanism plays an important role in the regulation of neurogenesis, differentiation and brain development ^10^. Furthermore, epigenetic alterations have been hypothesized to contribute to neurodevelopmental disorders ^11^, including ADHD ^12^, ASD ^13,14^, or borderline personality disorder ^15^.

DNA methylation can be complementary if it involves both alleles or non-complementary when it affects only one allele, as in chromosome X inactivation in females or allele-specific methylation (ASM) ^6^. ASM is a common mechanism by which single nucleotide variants determine differential methylation levels of CpG sites. ASM can alter promoter activity, leading to allele-specific expression ^16^ in combination with other still quite unknown factors, such as environmental effects ^6^. It is quantitative and heterogeneous across tissues and individuals ^6^. The environment affects DNA methylation leading to changes in gene regulation, although the underlying mechanism is still not well understood ^17^. It has been suggested that, during embryonic development, ASM regions could be especially sensitive to environmental effects ^6^. Investigating SNPs that display allele-specific methylation could help identify risk variants for common diseases, including neuropsychiatric disorders ^18^, as shown by recent studies of BD and schizophrenia ^9,19^.

The present study investigated the possible contribution of ASM to ADHD using data from the largest GWAS meta-analysis performed to date in ADHD ^4^. We assessed its possible effect on gene expression and on brain volumes to identify new genes contributing to the disorder.

## MATERIALS AND METHODS

### Selection of Allele-Specific Methylation SNPs

The SNP selection was made based on the results of two previous studies ^20,21^, which identified ASM variants in multiple brain regions of *post-mortem* human samples. Gibbs *et al.*, 2010 considered four brain regions (cerebellum, frontal cortex, caudal pons and temporal cortex) of 150 subjects and Zhang *et al.*, 2010 used only the cerebellum of 153 subjects.

In the study by Zhang *et al.*, 2010 a total of 12,117 SNP-CpG pairs associations were reported in cerebellum, and Gibbs *et al.*, 2010 listed a total of 12,135 SNP-CpG pairs in frontal cortex, 11,374 in caudal pons, 16,734 in temporal cortex and 12,102 in cerebellum (Fig. 1). We combined the information from both studies and obtained a total of 43,132 SNP-CpG pairs involving 33,944 different SNPs and 5,306 CpG sites (Fig. 1). We considered all the ASM SNPs from all the tissues in the two studies, as there are multiple SNP-CpG pairs in common between them (Fig. S1).

**Fig 1.**
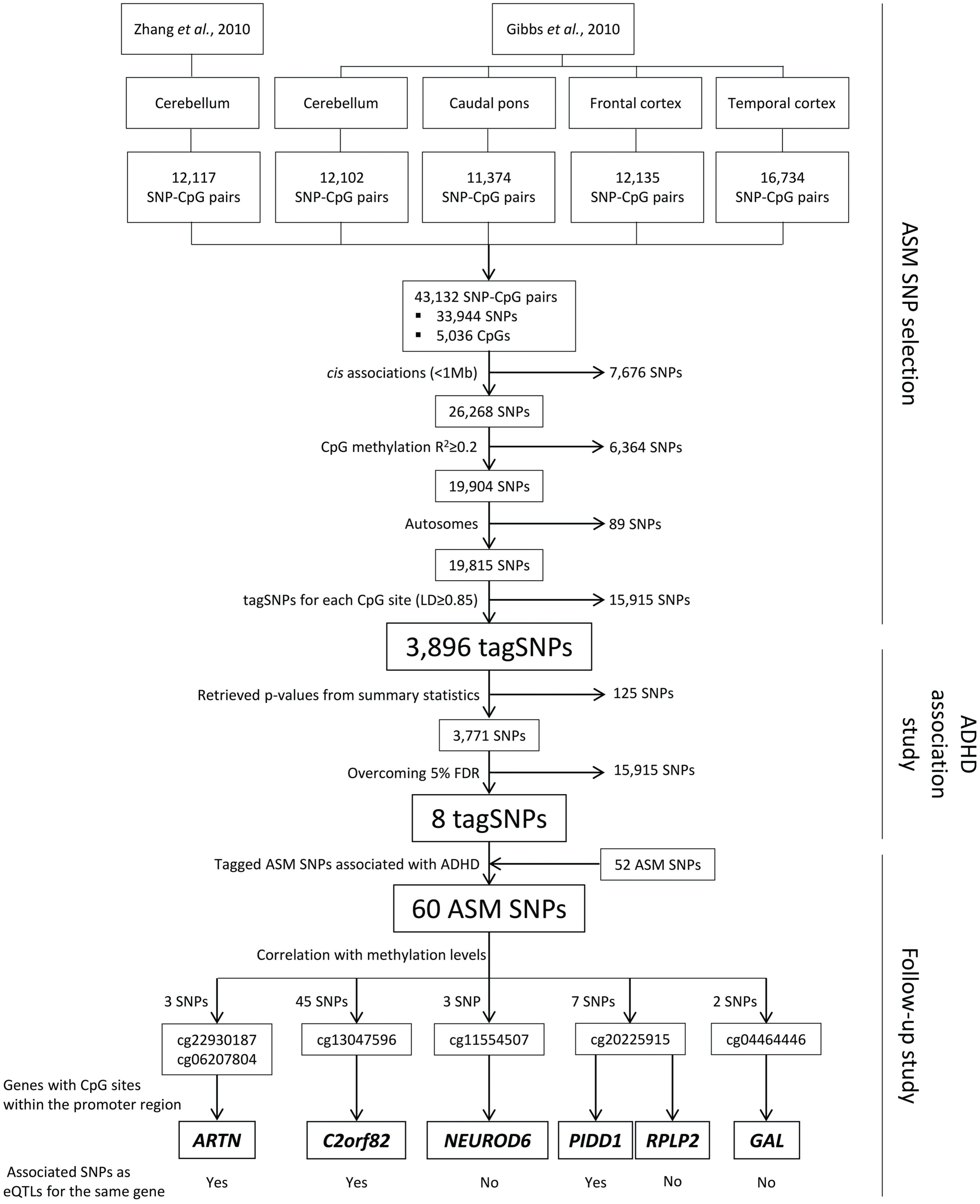
Selection of allele-specific methylation (ASM) SNPs from two previous studies and association results obtained for ASM variants in ADHD. SNPs tested in the ADHD GWAS meta-analysis and multiple testing correction. SNPs correlating with differential methylation of CpG sites and eQTLs in brain regions (only for genes in which the CpG site lies < 5 kb from the transcription start site) are depicted.

We subsequently applied different filters to generate a sub-list of 3,896 SNPs out of these 33,944 variants to minimize redundancy: associations in *cis* between the SNP and the CpG site, correlation of the SNP with methylation levels of the CpG (R^2^) ≥ 0.2, as performed in Gibbs *et al.* 2010 ^20^. We considered only autosomal SNPs and selected tagSNPs for each CpG site (r^2^ ≥ 0.85), by assessing linkage disequilibrium (LD) with Haploview software ^22^ using the Central European (CEU) reference panel from 1000 Genomes Project Phase 3 ^23^ (Fig. 1).

### Case-control GWAS datasets

We explored the selected ASM SNPs in the summary statistics from a meta-analysis of 11 independent GWAS of ADHD conducted by the Psychiatric Genomics Consortium (PGC) and iPSYCH. This case-control study investigated 8,047,420 markers in 20,183 cases and 35,191 controls from Europe, USA, Canada, and China, with patients diagnosed according to the criteria detailed in Demontis *et al.*, 2017 ^4^.

### Statistical analysis

From our initial selection of 3,896 ASM SNPs, information of 3,771 SNPs could be retrieved (96.8%) from the summary statistics of the ADHD GWAS meta-analysis ^4^. False Discovery Rate (FDR) was applied to correct for multiple testing. We used the q-value package for R ^24^ and obtained a threshold p-value of 6.78e-05 corresponding to a 5% FDR. CpG sites highlighted by SNPs that were significant at this FDR threshold were followed-up in further analyses (Fig. 1). We also performed Bonferroni correction for multiple testing (p ≤ 1.32e-05; 0.05/3771 SNPs) although it was too stringent in our study as the considered SNPs are not completely independent. Finally, we considered and retrieved p-values of those SNPs in high LD (r^2^ ≥ 0.85) with the previous ones that correlate in *cis* with the methylation levels of the same CpG sites (R^2^ ≥ 0.2).

### Functional annotation of associated ASM SNPs

We applied four methods to obtain information about the possible functional impact of the ASM SNPs that were associated with ADHD. First, we evaluated the presence of possible enhancer or promoter regions using the Haploreg v4.1 tool ^25^. To do this, we considered histone modifications related to enhancer regions (H3K4me1 and H3K27ac) and promoters (H3K4me3 and H3K9ac) of 10 different brain regions (hippocampus middle, substantia nigra, anterior caudate, cingulate gyrus, inferior temporal lobe, angular gyrus, dorsolateral prefrontal cortex, germinal matrix and male and female fetal brain). Second, we evaluated the effect on gene expression through an eQTL analysis using GTEx data (Release V7) ^26^. We considered eQTL information for all available brain tissues: amygdala, anterior cingulate cortex (BA24), caudate basal ganglia, cerebellar hemisphere, cerebellum, cortex, frontal cortex (BA9), hippocampus, hypothalamus, nucleus accumbens basal ganglia, putamen basal ganglia, spinal cord cervical c-1 and substantia nigra. Third, we considered all the SNPs, not only ASMs, located within ± 1Mb from the transcription start site (TSS) of each gene to infer if the genetically determined expressions of genes of interest correlated with ADHD. This analysis was carried out using MetaXcan ^27^, the input being the summary statistics of the ADHD GWAS meta-analysis ^4^ and prediction models trained with RNA-Seq data of 10 GTEx ^26^ brain tissues and CommonMind ^28^ dorsolateral prefrontal cortex. The SNP covariance matrices were generated using the 1000 Genomes Project Phase 3 ^23^ EUR genotypes of the prediction model SNPs. Bonferroni correction for multiple testing was considered (p ≤ 2.27e-03; 0.05/22 SNPs). Finally, we examined the possible influence of our identified variants on subcortical brain structures. We obtained the summary statistics of a GWAS meta-analysis of eight MRI volumetric measures (nucleus accumbens, amygdala, caudate nucleus, hippocampus, pallidum, putamen, thalamus and intracranial volume (ICV)) produced by the Enhancing Neuro Imaging Genetics through Meta-Analysis (ENIGMA) consortium ^29^. This ENIGMA2 discovery sample included 13,171 subjects of European ancestry and contained association results between seven million markers and variance in the volumes of the mentioned structures ^29^; we applied the Bonferroni correction (p ≤ 1e-03; 0.05/50 SNPs)

## RESULTS

We investigated the possible association with ADHD of SNPs that show allele-specific methylation in brain regions. Starting from two previous studies ^20,21^ that describe ASM in brain tissues we obtained 43,132 SNP-CpG pairs involving 33,944 SNPs and 5,306 CpG sites (Fig. 1). We observed some overlaps and redundancies between studies and tissues (Fig. S1), so we performed a selection process ending up with a list of 3,896 ASM tagSNPs (Fig. 1). Eight ASM tagSNPs were significantly associated with ADHD after correcting for multiple comparisons (5% FDR, p ≤ 6.78e-05). These eight SNPs correlated with differential methylation at six CpG sites in cis (three for cg20225915, two for both cg22930187 and cg06207804, and one for each of cg13047596, cg11554507 and cg04464446) in different brain areas (Figs. 1 to 4). Although the explored SNPs are not completely independent in terms of linkage disequilibrium (LD), we also applied the Bonferroni correction (p ≤ 1.32e-05; 0.05/3771 SNPs) and three of the eight ASM tagSNPs remained statistically significant, all correlating with differential methylation at the cg20225915 site (Table 1). As considering only tagSNPs may overlook true causal variants, we retrieved association results from all the 52 ASM SNPs tagged by the previous ones (LD; r^2^ ≥ 0.85), ending up with 60 variants in eight LD blocks that show association with ADHD and correlate with methylation levels at six CpG sites (Table S1). Regional association plots of the eight tagSNPs can be found in Figs. S2 to S9 and LD patterns among the 60 variants are described in Figs. S10 to S14.

**Table 1.** Allele-Specific Methylation (ASM) tagSNPs associated with ADHD.

| SNP | Chr | Pos | Alleles |  | OR | Freq A1 |  | Meta-analysis p |
| --- | --- | --- | --- | --- | --- | --- | --- | --- |
|  |  |  | A1 | A2 |  | Cases | Controls |  |
| rs2906458 | 1 | 44336389 | A | <u>G</u> | 0.94 | 0.74 | 0.756 | 3.01E-05 |
| rs7412307 | 1 | 44433864 | <u>C</u> | G | 1.07 | 0.185 | 0.172 | 2.82E-05 |
| rs11676216 | 2 | 233706368 | T | <u>C</u> | 0.95 | 0.646 | 0.654 | 6.78E-05 |
| rs4140961 | 7 | 31349352 | <u>A</u> | G | 1.06 | 0.597 | 0.592 | 6.05E-05 |
| **rs7104929** | **11** | **784340** | **C** | **<u>G</u>** | **0.94** | **0.512** | **0.526** | **7.89E-06** |
| **rs7479101** | **11** | **802115** | **A** | **<u>G</u>** | **0.93** | **0.317** | **0.33** | **5.90E-06** |
| **rs4131364** | **11** | **812188** | **<u>A</u>** | **G** | **1.07** | **0.517** | **0.502** | **1.60E-06** |
| rs11600377 | 11 | 68785803 | <u>A</u> | G | 1.06 | 0.731 | 0.72 | 4.38E-05 |

SNP: Single Nucleotide Polymorphism; Chr: Chromosome; Pos: Position (build hg19); A1: Allele 1; A2: Allele 2; OR: Odds Ratio (calculated on A1); Freq A1: Frequency of allele 1; In bold: Significant tagSNPs overcoming Bonferroni correction for multiple testing; Underlined allele: risk allele for ADHD; meta-analysis p: p-value obtained from the PGC+iPSYCH meta-analysis (Demontis *et al.*, 2017).

The ADHD risk alleles of the 60 SNPs and their effects on CpG methylation levels are reported in Figs. 2 to 4 and Table S1. Consistently, the direction of the effect of the risk alleles on methylation levels is the same for all the SNPs in each LD block, correlating with the same CpG site. Thus, the risk alleles correlate with decreased methylation of cg22930187, cg06207804, cg11554507 and cg20225915 and with increased methylation of cg13047596 and cg04464446 ^20,21^ (Figs. 2, 3, 4 and Table S1).

**Fig 2.**
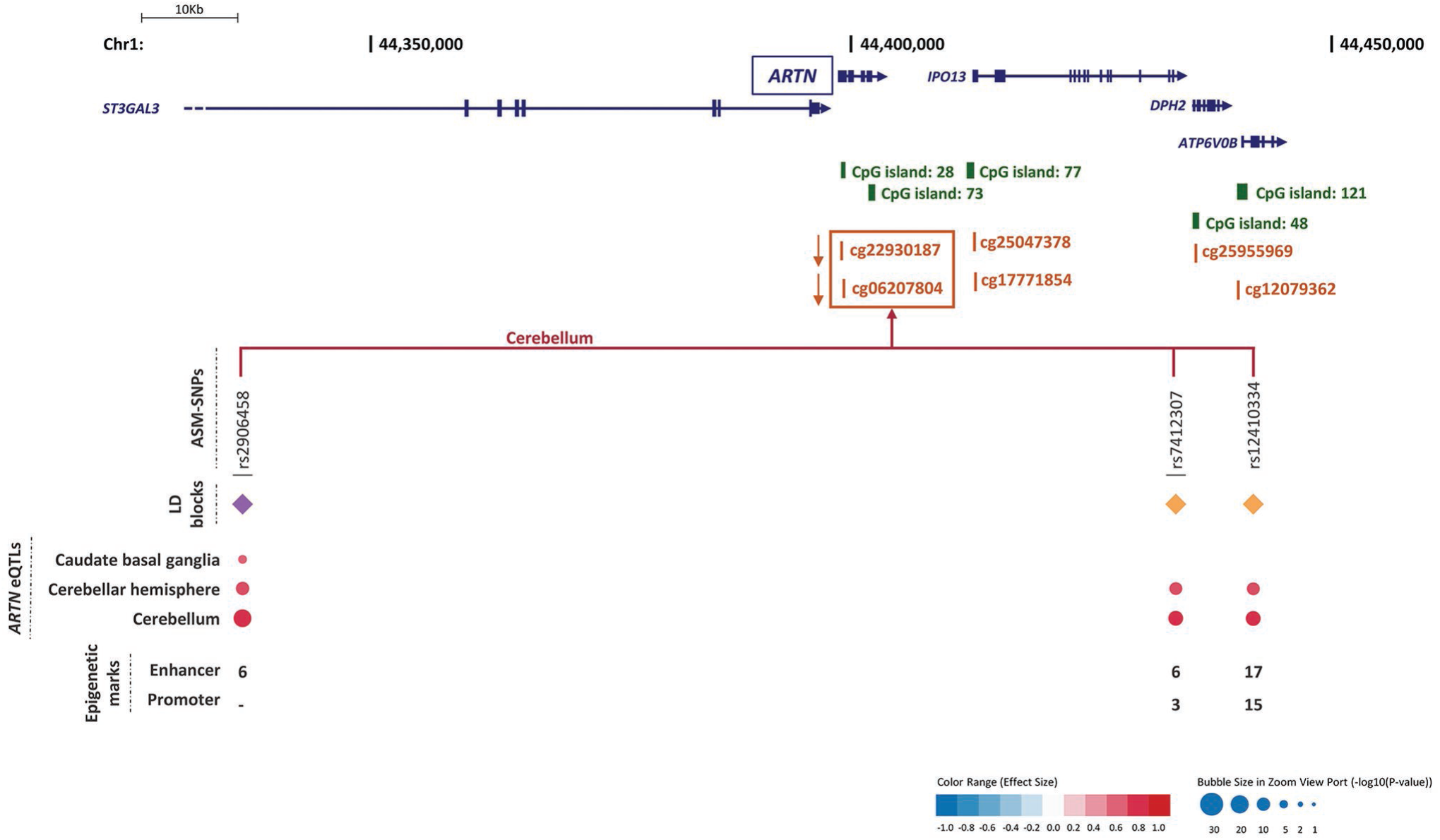
Genomic context of ASM variants, and methylation and eQTL information for cg22930187 and cg06207804. Genes are depicted in dark blue, showing the direction of transcription with an arrow; CpG sites inspected in the reference studies appear in brown; framed CpG sites indicate those sites showing differential levels of methylation for the associated ASM SNPs, and brown arrows indicate the effect on methylation of the ADHD risk variants, with indication of the brain regions where the ASMs were described. The tagSNPs are underscored. The colored rhombuses show the LD blocks present in each region. The colored dots for eQTLs indicate the effect on gene expression of the ADHD risk allele, according to the legend (red: over-expression, blue: under-expression). The number of enhancer (H3K4me1 and H3K27ac) and promoter (H3K4me3 and H3K9ac) histone marks found in the different brain areas are displayed for each SNP. ‘-’ indicates no known enhancer or promoter histone marks.

**Fig 3.**
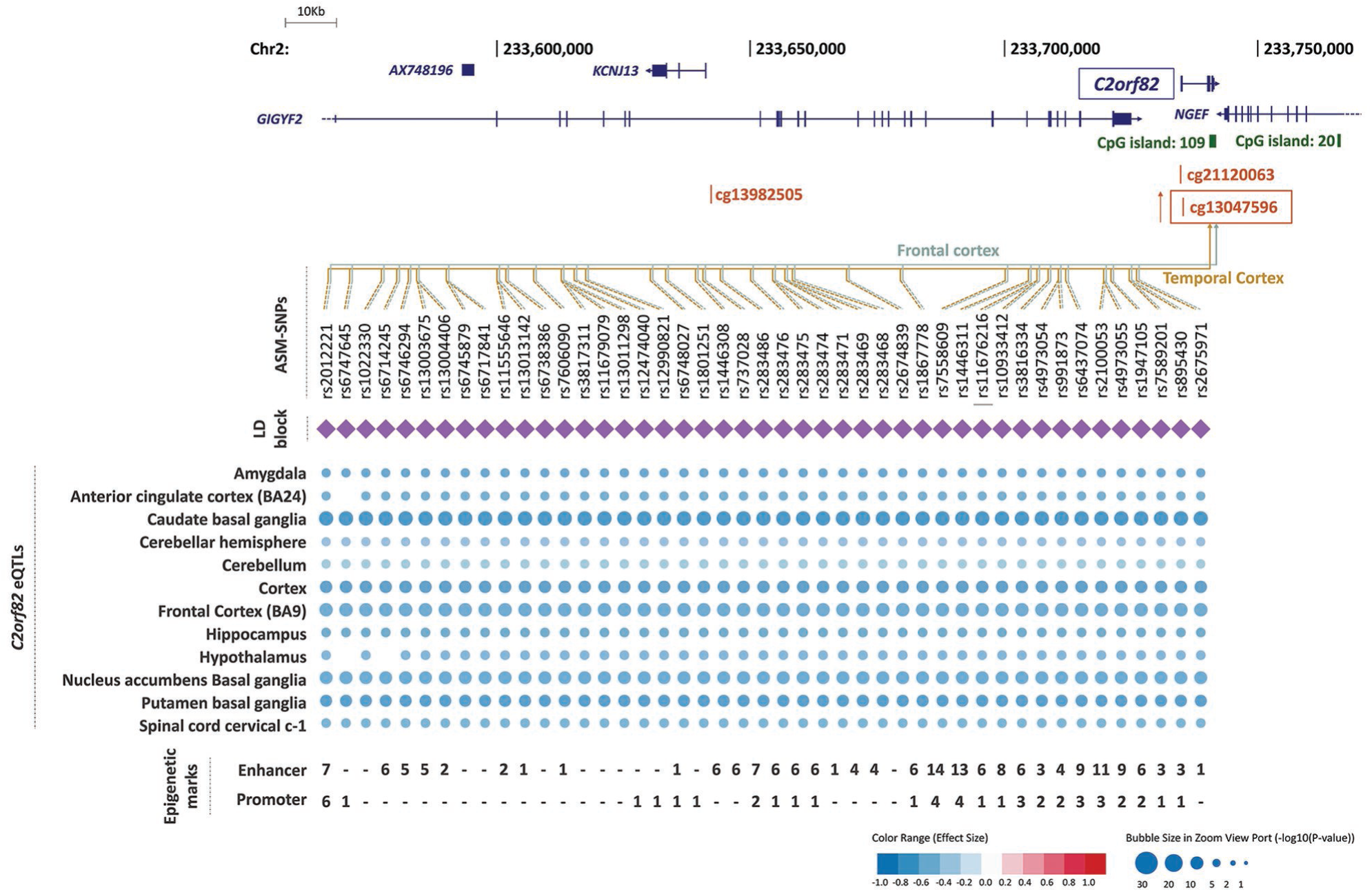
Genomic context of ASM variants, and methylation and eQTL information for cg13047596. Genes are depicted in dark blue, showing the direction of transcription with an arrow; CpG sites inspected in the reference studies appear in brown; framed CpG sites indicate those sites showing differential levels of methylation for the associated ASM SNPs, and brown arrows indicate the effect on methylation of the ADHD risk variants, with indication of the brain regions where the ASMs were described. The tagSNPs are underscored. The colored rhombuses show the different LD blocks present in each region. The colored dots for eQTLs indicate the effect on gene expression of the ADHD risk allele, according to the legend (red: over-expression, blue: under-expression). The number of enhancer (H3K4me1 and H3K27ac) and promoter (H3K4me3 and H3K9ac) histone marks found in the different brain areas are displayed for each SNP. ‘-’ indicates no known enhancer or promoter histone marks.

**Fig 4.**
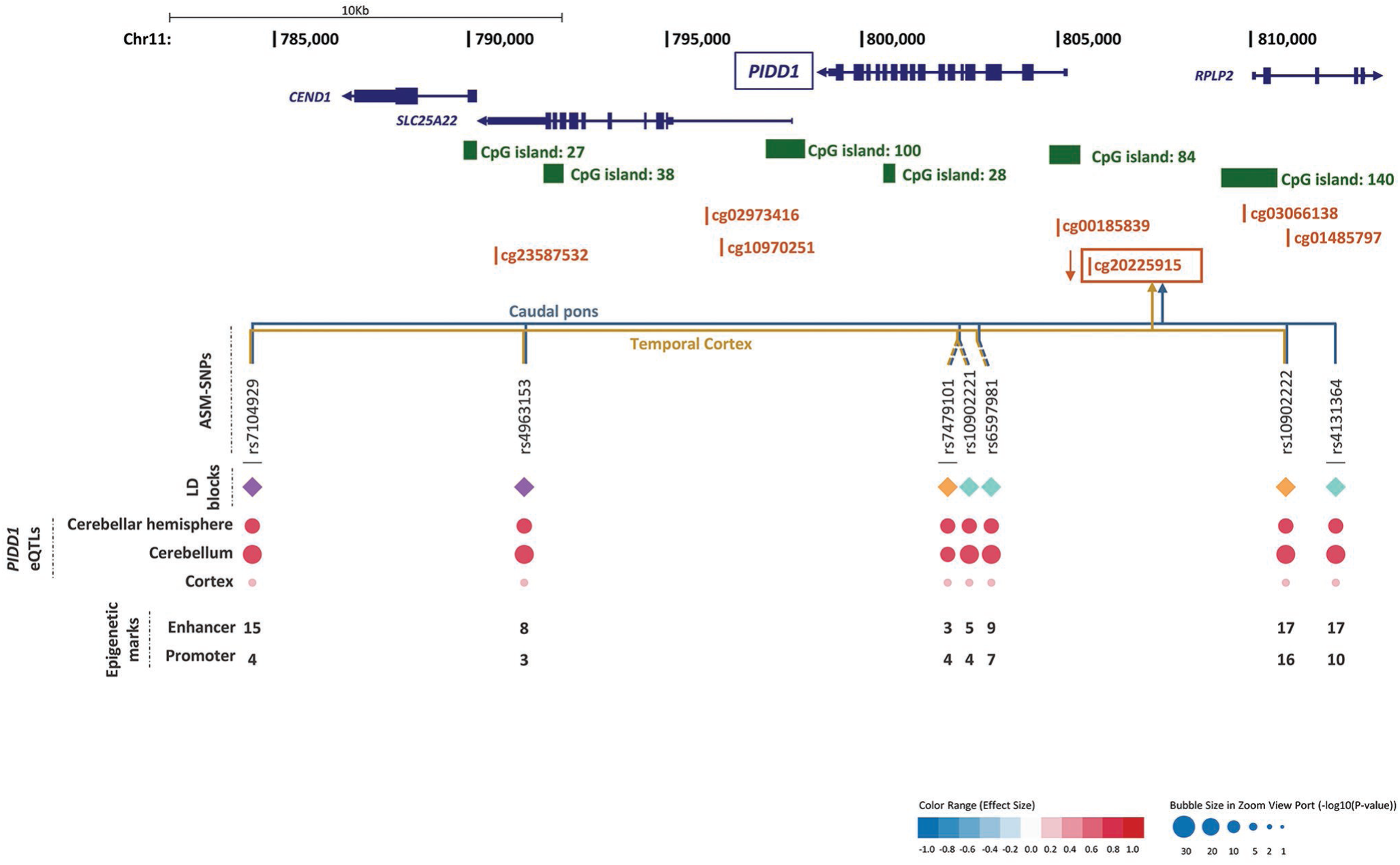
Genomic context of ASM variants, and methylation and eQTL information for cg20225915. Genes are depicted in dark blue, showing the direction of transcription with an arrow; CpG sites inspected in the reference studies appear in brown; framed CpG sites indicate those sites showing differential levels of methylation for the associated ASM SNPs, and brown arrows indicate the effect on methylation of the ADHD risk variants, with indication of the brain regions where the ASMs were described. The tagSNPs are underscored. The colored rhombuses show the LD blocks present in each region. The colored dots for eQTLs indicate the effect on gene expression of the ADHD risk allele, according to the legend (red: over-expression, blue: under-expression). The number of enhancer (H3K4me1 and H3K27ac) and promoter (H3K4me3 and H3K9ac) histone marks found in the different brain areas are displayed for each SNP. ‘-’ indicates no known enhancer or promoter histone marks.

All six identified CpG sites are located in possible promoter regions (less than 5,000 bp upstream from a TSS) of six genes: *ARTN* (cg22930187 and cg06207804), *C2orf82* (cg13047596), *NEUROD6* (cg11554507), *PIDD1* (cg20225915), *RPLP2* (cg20225915) and *GAL* (cg04464446), all of them expressed in brain. Furthermore, most of the ASM SNPs have histone marks in different brain areas related to enhancer or promoter regions (Figs. 2 to 4 and Tables S2 to S6).

We subsequently assessed the possible effect of those 60 SNPs on gene expression and observed that all the SNPs from six LD blocks are eQTLS for three of the six genes, where the differentially methylated CpG sites lie in the possible promoter regions: *ARTN, C2orf82* and *PIDD1* (Figs. 2 to 4 and Table S1). The ADHD risk alleles are associated with increased expression of *ARTN* (in caudate basal ganglia, cerebellar hemisphere and cerebellum) and *PIDD1* (in cerebellar hemisphere, cerebellum and cortex) and with decreased expression of *C2orf82* (in amygdala, anterior cingulate cortex, caudate basal ganglia, cerebellar hemisphere, cerebellum, cortex, frontal cortex, hippocampus, hypothalamus, nucleus accumbens basal ganglia, putamen basal ganglia and spinal cord cervical c-1) (Figs. 2 to 4).

Consistently, all risk variants for ADHD showed opposite directions for methylation and gene expression levels, as expected for CpG sites lying in promoter regions (Table S1). Also, all the ADHD risk alleles from different LD blocks within each gene contribute in the same direction to its methylation and to its expression in all the investigated tissues (Figs. 2 to 4 and Table S1).

Furthermore, we assessed whether the effect on gene expression predicted for these genes correlate with ADHD by considering all variants within ±1 MB from the TSS (and not only the ASM SNPs) using MetaXcan. We found significant associations of gene expression with ADHD for the same three genes in multiple brain tissues: *ARTN, PIDD1* showed increased expression (3.57 < Z-score < 4.19 and 3.57 < Z-score < 5.37 respectively) and *C2orf82* with a decreased expression (−3.64 < Z-score < −3.07) (Table S7), all of them surviving the Bonferroni correction for multiple testing (p ≤ 2.27e-03). Consistently, for these genes, the predicted direction of the effects for the gene expression changes according to MetaXcan are the same as the ones observed in the eQTL analysis.

We also evaluated the correlation of the 60 ADHD-associated SNPs with subcortical brain volume changes in ENIGMA2 data, where we could retrieve information for 50 SNPs. We found that most the SNPs from three LD blocks are nominally significant in at least one of these three subcortical brain regions: accumbens, caudate and thalamus (Table S8). Most of the SNPs correlating with cg13047596 (near *C2orf82*) and with cg04464446 (near *GAL*), correlate with both accumbens and caudate volumes, while the only SNP correlating with cg11554507 (near *NEUROD6*) present in ENIGMA2 correlates with thalamus volume (Table S8).

## DISCUSSION

This study is the first comprehensive assessment of the contribution to ADHD of genetic variants altering methylation in brain. We identified a total of 60 variants from eight LD blocks associated with ADHD that correlated with differential levels of methylation at six different CpG sites ^20,21^ (Tables 1 and S1). All the variants from six out of the eight LD blocks alter the methylation of CpG sites lying in the possible promoter regions and are also eQTLs for one of the following three genes in brain: *ARTN, C2orf82* and *PIDD1* (Figs. 2 to 4 and Table S1). It is well known that DNA methylation in promoter regions inversely correlates with levels of gene expression ^17^, and all the ASM variants that we found associated with ADHD in our study are concordant with this statement.

The *ARTN* gene, highlighted by two tagSNPs, encodes Artemin, a ligand of the *GDNF* family (glial cell line-derived neurotrophic factor). Artemin supports the survival of sensory and sympathetic peripheral neurons in culture by interacting with GFRα3-RET and possibly also of dopaminergic neurons of the ventral mid-brain through activation of GFRα1-RET complex ^30^. Gene Ontology (GO) pathways link it to key neurodevelopmental functions: axon guidance (GO:0007411), neuroblast proliferation (GO:0007405) and peripheral nervous system development (GO:0007422). Risk alleles for ADHD lead to an overexpression of *ARTN*. Previously, overexpression of *ARTN* has been studied in transgenic mice and been linked to an increase of neuron excitability that leads to hypersensitivity ^31,32^. Another study in *ARTN* knockout mice reported aberrations in the sympathetic nervous system related to migration and axonal projection ^33^. The *C2orf82* gene (also known as *SNORC*) was highlighted by one tagSNP and it encodes a proteoglycan transmembrane protein that is expressed in brain more than in other tissues ^26^. Little is known about its function. Finally, *PIDD1* was highlighted by three tagSNPs. It is a cell life regulator gene and it has been linked to apoptotic and anti-apoptotic pathways. The PIDD protein initiates apoptosis as a component of the PIDDosome together with RAIDD (RIP-associated ICH-1/ECD3-homologous protein with a death domain) and procaspase-2 ^34^ and it also activates an anti-apoptotic pathway involving the transcription factor NF-κB in response to genotoxic stress ^35^

The ASM SNPs from six LD blocks associated with ADHD are eQTLs for one of the following genes: *ARTN, C2orf82* and *PIDD1* in several brain regions (Figs. 2 to 4). Also, when we considered all the variants in these genes (±1 MB from the TSS), *ARTN, C2orf82* and *PIDD1* were also predicted to be differentially expressed in the same direction in ADHD by MetaXcan (Table S7). The cerebellum is an expression site common to all these eQTLs, which can be explained in part by the fact that the initial SNP selection is enriched in ASM SNPs of this brain region (Fig. S1). Structural anomalies in cerebellum have been reported in ADHD individuals through neuroimaging studies ^36–38^. ADHD is suggested to be related to problems in processing temporal information, and cerebellum, basal ganglia and prefrontal cortex are involved in the temporal information processing ^39^. Cerebellar developmental trajectories and hippocampal volumes are linked to the severity of ADHD symptoms ^40–42^. Structural and functional abnormalities in cerebellum and basal ganglia have further been associated with motor impairments ^43^, which are frequent in nearly half of ADHD cases ^44^. Other areas identified through our eQTL analysis have also been related to ADHD, for instance: i) Remarkable differences in the shapes of caudate-putamen basal ganglia and smaller volumes have been reported in ADHD boys in comparison to typically developing individuals ^45–48^; ii) in adult males with ADHD, right caudate volume correlates with poor accuracy on sensory selection tasks ^49^ and also with hyperactivity/impulsivity ^50,51^; iii) nucleus accumbens is one of the areas, together with the caudate nucleus, putamen, amygdala, and hippocampus, reported to be structurally altered in the brain of ADHD patients ^52^. A delayed cortical development e.g. in prefrontal regions has been linked to ADHD ^53,54^ with below median intelligence quotient (IQ) ^55^, with the exception of primary motor cortex which shows slightly earlier maturation ^54^; and iv) there is speculation about the role of cortical thickness ^56^, cortical volume ^57^ and functional connectivity in the anterior cingulate cortex ^58^, the region for cognitive control, attention, affect and drive ^59^ in ADHD ^59–62^. These fronto-subcortical structures (lateral prefrontal cortex, dorsal anterior cingulate cortex, caudate, and putamen) and pathways are rich in catecholamines which are involved in the pharmacological treatments for ADHD ^44,48,53,63^. Remarkably, most of the ASM SNPs in the LD block for *C2orf82*, also nominally correlate with increased volumes of accumbens and caudate subcortical regions (Table S8). It is interesting that these SNPs have the higher effect sizes as eQTLs in caudate basal ganglia (Fig. 3), one of the subcortical regions which volume correlates with genotype variation at these SNPs.

Interestingly, the methylation of cg20225915 has also been associated with *PIDD1* expression in peripheral blood ^64^, turning it into a good candidate as a biomarker. Also, two of the three highlighted genes had previously been related to other psychiatric diseases. The expression of *ARTN and C2orf82* was found to be altered in blood of MDD ^65^ and schizophrenia patients ^66,67^, respectively. The fact that these genes have previously been related to neuropsychiatric disorders that are often comorbid with ADHD ^68^ make them appealing candidates to be pursued.

*ARTN* is the only gene highlighted in our study that is present in one of the top regions reported in the ADHD GWAS meta-analysis ^4^, although it did not contain SNPs surviving genome-wide significance. The GWAS findings in the region could be accounted for by one of several genes: *ST3GAL3, PTPRF, KDM4A, RP11-184I16.4, XR_246316.1, KDM4A-AS1* and *SLC6A9*. *ST3GAL3* had the most signals. Although two of our ASM variants associated with ADHD are intronic to *ST3GAL3*, this gene was not highlighted in our study as none of the associated variants correlated with differential methylation of CpG sites near the *ST3GAL3* transcription start site (distance from the nearest CpG site: 197 Kb) or were eQTLs for the gene in brain tissues. Instead, these SNPs correlated with a nearby gene, *ARTN*, both in terms of methylation and gene expression. This suggests the importance of finding functional connections between disease-associated SNPs and genes, besides considering the genes in the physical vicinity of variants. Furthermore, another of our highlighted genes, *PIDD1*, although not being among the top findings in the ADHD GWAS meta-analysis ^4^, it is pointed out by the gene-based association analysis performed in the same study.

Genetic variants surpassing genome-wide significance in GWAS explain only a small part of the SNP-based heritability and associations not reaching the significance threshold also contribute to disease susceptibility ^4,8^. An omnigenic model has been recently proposed suggesting that these subthreshold variants could point at regulatory elements of core genes ^7^. Indeed, a previous study on a cardiovascular cardiac phenotype reported that nominally significant associations are enriched in enhancer regions ^8^. Therefore, although none of the variants that we identified in our study display genome-wide significant association with ADHD, they may contribute to the susceptibility to ADHD, as they do have a functional impact (methylation, expression and in some cases brain structure) on genes that are expressed in brain. It is also supported by the data shown in Tables S2, S3, S4, S5 and S6, showing that most of our SNPs are related to different histone marks linked to possible promoter or enhancer regions. Brain-specific ASM information has also been utilized to detect key genes and pathways in bipolar disorder ^19^. Also, enrichment of brain ASM in a schizophrenia GWAS in comparison to non-psychiatric GWAS reinforces the rationale of utilizing ASM SNPs to highlight genes that are relevant to psychiatric disorders from GWAS data ^9^. None of our highlighted SNPs or CpG sites overlap with the ones described in a previous study relating ASM to schizophrenia ^69^.

There are some strengths and limitations in our study that should be discussed Strengths: i) We used the largest meta-analysis of ADHD GWAS performed so far, including around 20,000 cases and 35,000 controls. ii) Two of the highlighted genes, *ARTN* and *C2orf82,* had previously been associated with other psychiatric disorders. iii) For two of the genes there is more than one LD block showing the same effect on CpG site methylation. iv) Our results are concordant with eQTL information that had been assessed in an independent sample, with all the SNP showing the opposite effect on methylation of the promoter region and on the expression of a given gene in brain (more promoter methylation and less gene expression or vice versa), even for the different LD blocks included in each region. Limitations: i) We did not perform a follow-up study to replicate our association findings in an independent sample. ii) The previous studies that we used for the selection of ASM SNPs where performed on different genotyping platforms that do not include all the existing SNPs in the genome, and therefore we could not test all possible ASMs. iii) We only considered cis-associated ASM variants, which are the vast majority, although non-cis ASM also occurs. iv) There is an overrepresentation of ASM SNPs from cerebellum compared to the other studied tissues.

To conclude, our study points to the *ARTN, C2orf82* and *PIDD1* genes as potential contributors to ADHD susceptibility. The identified risk variants regulate the methylation levels of different CpG sites located in promoter regions and they inversely correlate with expression of the corresponding genes in brain. This finding is supported by a prediction of increased expression of *ARTN* and *PIDD1* and a decreased expression of *C2orf82* in ADHD. Moreover, most of the variants correlating with methylation at cg13047596 (near *C2orf82*) have an influence on accumbens and caudate volumes. Further studies are required to elucidate the mechanisms by which these genes contribute to ADHD.

## ACKNOWLEDGMENTS

Major financial support for this research was received by BC from the Spanish ‘Ministerio de Economía y Competitividad’ (SAF2015-68341-R) and AGAUR, ‘Generalitat de Catalunya’ (2017-SGR-738). The research leading to these results has also received funding from the European Union Seventh Framework Programme [FP7/2007-2013] under grant agreement n° 602805 and from the European Union H2020 Program [H2020/2014-2020] under grant agreements n° 667302 and n° 643051, the latter supporting the contract of AS. LP-C and JC-D were supported by ‘Generalitat de Catalunya’ (2016 FI_B 00728 and 2015 FI_B 00448, respectively). LP-C was also supported by ‘Ministerio de Educación, Cultura y Deporte’ (FPU15/03867). AS was supported by the European Union H2020 Program [H2020/2014-2020] under grant agreements n° 643051 by the NF-C was supported by contracts of the ‘Centro de Investigación Biomédica en Red de Enfermedades Raras’ (CIBERER). VR was supported by the Graduate School of Health from the Univeristy of Aarhus. The iPSYCH team acknowledges support from the Lundbeck Foundation. Finally, SF was supported by the European Union’s Seventh Framework Programme for research, technological development and demonstration under grant agreement no 602805, the European Union’s Horizon 2020 research and innovation programme under grant agreements n° 667302 and n° 728018 and NIMH grants 5R01MH101519 and U01 MH109536-01.

We are thankful to Roser Corominas (Universitat de Barcelona, Barcelona) for helpful advice. We are also grateful to the ADHD Working Group of the Psychiatric Genomics Consortium (PGC) and the iPSYCH team for distributing the summary statistics of the ADHD GWAS meta-analysis. Access to the PGC ADHD data was obtained through dbGaP project number 10608 that includes the following datasets: phs000016.v2.p2, phs000407.v1.p1, phs000358.v1.p1 and phs000490.v1.p1. We thank the ENIGMA consortium for sharing the summary statistics of genome-wide association meta-analyses of MRI phenotypes.

